# *delilah, prospero* and *D-Pax2* constitute a gene regulatory network essential for the development of functional proprioceptors

**DOI:** 10.1101/2021.04.07.438778

**Authors:** Adel Avetisyan, Yael Glatt, Maya Cohen, Yael Timerman, Nitay Aspis, Atalya Nachman, Naomi Halachmi, Ella Preger-Ben Noon, Adi Salzberg

## Abstract

Coordinated animal locomotion depends on the development of functional proprioceptors. While early cell-fate determination processes are well characterized, little is known about the terminal differentiation of cells within the proprioceptive lineage and the genetic networks that control them. In this work we describe a gene regulatory network consisting of three transcription factors-Prospero (Pros), D-Pax2 and Delilah (Dei)-that dictates two alternative differentiation programs within the proprioceptive lineage in Drosophila. We show that D-Pax2 and Pros control the differentiation of cap versus scolopale cells in the chordotonal organ lineage by, respectively, activating and repressing the transcription of *dei.* Normally, D-Pax2 activates the expression of *dei* in the cap cell but is unable to do so in the scolopale cell where Pros is co-expressed. We further show that D-Pax2 and Pros exert their effects on *dei* transcription via a 262 bp chordotonal-specific enhancer in which two D-Pax2- and three Pros-binding sites were identified experimentally. When this enhancer was removed from the fly genome, the cap- and ligament-specific expression of *dei* was lost, resulting in loss of chordotonal organ functionality and defective larval locomotion. Thus, coordinated larval locomotion depends on the activity of a *dei* enhancer that integrates both activating and repressive inputs for the generation of a functional proprioceptive organ.

## INTRODUCTION

A central question in developmental biology is how different cells that originate in the same lineage and develop within the same organ, acquire unique identities, properties and specialized morphologies. One of the common mechanisms involved in cell fate diversification within a cell lineage is asymmetric cell division in which cytoplasmic determinants of the mother cell differentially segregate into one of the two daughter cells. This asymmetry is then translated into differential gene expression and the activation of cell type-specific gene regulatory networks (GRN) that dictate the differentiation programs of cells with unique properties. The transition from a primary cell fate to the characteristic phenotype of a fully differentiated cell involves complex GRNs in which numerous genes regulate each other’s expression. Despite this complexity, genetic analyses in well-characterized developmental systems can often reveal elementary interactions in small GRNs which dictate a specific cell fate, or a specific feature of the differentiating cell.

Many of the core components and the central processes underlying asymmetric cell divisions and primary cell fate decisions have been uncovered in studies performed on the central and peripheral nervous system (PNS) of *Drosophila* (*e.g* Knoblich, 2008; Schweisguth, 2015). The PNS of Drosophila contains two classes of multicellular sensory organs, external sensory organs and chordotonal organs (ChOs), whose lineages share a similar pattern of asymmetric cell divisions (Lai and Orgogozo, 2004). In both types of organs, the neuron and support cells, which collectively comprise the sensory organ, arise from a single sensory organ precursor cell (SOP) through a sequence of precisely choreographed asymmetric cell divisions (Reeves and Posakony, 2005). Unlike the primary cell-fate specification, which has been extensively investigated, the process of terminal differentiation of the post-mitotic progeny is poorly understood. We are using the larval lateral pentascolopidial ChO (LCh5) as a model system to study cell fate diversification within a sensory lineage.

The LCh5 organ is composed of five mechano-sensory units (scolopidia) that are attached to the cuticle via specialized epidermal attachment cells. Each of the five scolopidia originates in a single precursor cell that divides asymmetrically to generate five of the six cell types that construct the mature organ: the neuron, scolopale, ligament, cap and cap-attachment cell (Brewster and Bodmer, 1995)(Figure 1A-C). Three of the five cap-attachment cells are rapidly removed by apoptosis, leaving two cap-attachment cells that anchor the five cap cells to the epidermis (Avetisyan and Salzberg, 2019). Later in development, following the migration of the LCh5 organ from the dorsal to the lateral region of the segment, a single ligament-attachment cell is recruited from the epidermis to anchor the five ligament cells to the cuticle (Inbal et al., 2004). The mature LCh5 organ responds to mechanical stimuli generated by muscle contractions that lead to relative displacement of the attachment cells and the consequent shortening of the organ (Hassan et al., 2019).

**Figure 1.**
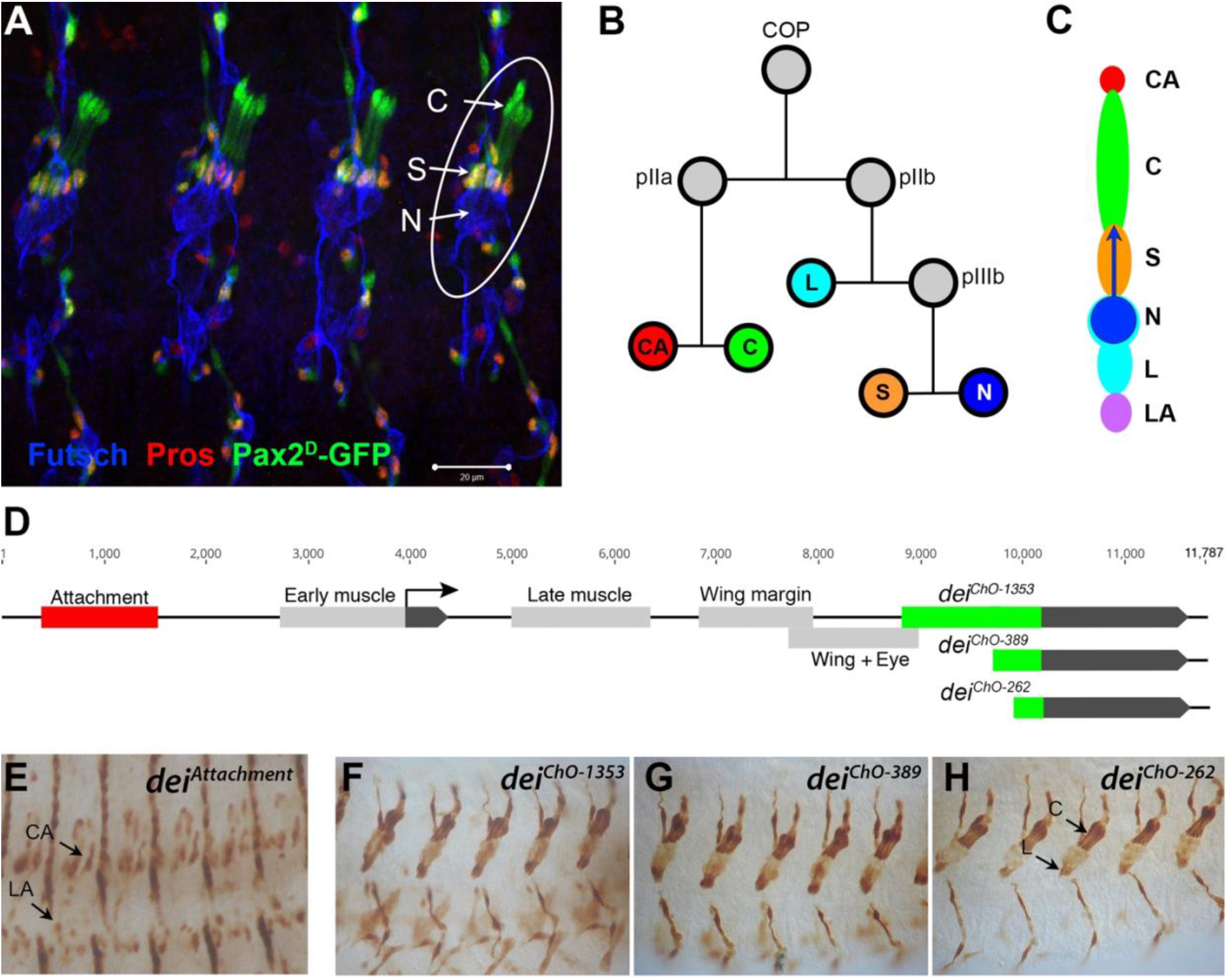
The LCh5 organ and the *dei* gene. (A) Four abdominal segments of a stage 16 embryo carrying the *sv/Pax2^*D1*^-GFP* reporter (green) which labels the cap and scolopale cells, stained with anti-Pros (red) which labels the nuclei of scolopale cells, and anti-Futsch (blue), which labels the neurons. One LCh5 organ is circled and the cap cells (C), scolopale cells (S) and neurons (N) are indicated. The cap-attachment, ligament-attachment and ligament cells are not stained. Scale bar = 20μm. (B-C) the ChO lineage (B) and schematic illustration of a single scolopidium (C). The neuron is depicted in blue, the scolopale cell in orange, the ligament cell in cerulean, the cap cell in green, the cap-attachment cell in red and the ligament-attachment cell in purple. (D) Schematic representation of the *dei* locus showing the two exons (black boxes) and the *cis* regulatory modules (CRM) that drive gene expression within the ChO lineage: the *dei*^*attachment*^ CRM which drives expression in the attachment cells (red box) and the *dei*^*ChO*^ CRM which drives expression in the cap and ligament cells (green box). The *dei*^*ChO*^ CRM was originally mapped to a 1353 bp fragment (*dei*^*ChO-1353*^) and was then delimited to a smaller 389 bp fragment located immediately upstream to the second exon (*dei*^*ChO-389*^). As part of this work, the ChO-specific CRM was further delimited to a 262 bp fragment (*dei*^*ChO-262*^). (E-H) The embryonic expression patterns driven by the *dei*^*attachment*^ (E), *dei*^*ChO-1353*^ (F), *dei*^*ChO-389*^ (G) and *dei*^*ChO-262*^ (H) enhancers are shown. C, cap cell; L, ligament cell; CA, cap-attachment cell, LA, ligament-attachment cell.

Very little is known about the unique cell type-specific differentiation programs that characterize each of the ChO cells, whose morphologies and mechanical properties differ dramatically from each other. To address this issue, we focus on the transcription factor Taxi wings/Delilah (Dei), an important regulator of cell adhesion (Egoz-Matia et al., 2011), which is expressed in the four accessory cell types (cap, ligament, cap-attachment and ligament-attachment) but is excluded from the neuron and the scolopale cell. Even though *dei* is expressed in all four accessory cells, its expression in these cells is differentially regulated. The transcription of *dei* in the ChO is controlled by two *cis*-regulatory modules (CRMs): The *dei*^*attachment*^ enhancer, located ~2.5Kb upstream of the *dei* transcription start site, drives expression in the cap-attachment and ligament-attachment cells (as well as tendon cells), whereas the *dei*^*ChO-1353*^ enhancer, an intronic 1353bp DNA fragment, drives *dei* expression specifically in the cap and ligament cells (Nachman et al., 2015) (Figure 1D-F). The *dei*^*attachment*^ enhancer was shown to be activated by the transcription factor Stripe, which is considered a key regulator of tendon cell development (Becker et al., 1997) and a known determinant of attachment cell identity (Klein et al., 2010). Here we provide a high-resolution dissection of the *dei*^*ChO-1353*^ enhancer and show that it integrates both activating and repressive cues to drive *dei* expression in cap and ligament cells while suppressing it in scolopale cells. We find that D-Pax2/Shaven (Sv), which is expressed in both branches of the cell lineage, is a positive regulator of *dei* and that Prospero (Pros) inhibits *dei* expression specifically in the scolopale cell. This small GRN is required for the realization of differentiation programs characterizing cap versus scolopale cell fates and is therefore essential for ChO functionality and coordinated larval locomotion.

## RESULTS

### Identifying D-Pax2/Sv as a putative direct activator of *dei* expression

To reveal the gene network that regulates *dei* expression in the ChO cells we used the ChO-specific *dei*^*ChO-1353*^ enhancer as an entry point. We first mapped the critical regulatory region within the *dei*^*ChO-1353*^ enhancer to a 389 bp fragment (*dei*^*ChO-389*^) that accurately recapitulated the expression pattern of the full *dei*^*ChO-1353*^ enhancer (Nachman et al., 2015) (Figure 1D, F-G). Subsequently, we narrowed down the critical regulatory region to an evolutionarily conserved 262 bp fragment *dei*^*ChO-262*^, which drives ChO-specific expression in a pattern indistinguishable from the expression pattern driven by the larger fragments *dei*^*ChO-1353*^ and *dei*^*ChO-389*^ (Figure 1D, F, H).

We combined two types of screens in an attempt to identify genes that regulate the expression of *dei* in the ChO lineage. The first screen was an RNAi-based phenotypic screen conducted in second instar larvae (Hassan et al., 2018). It capitalized on a reporter fly strain in which the cap and ligament cells expressed green fluorescent protein (GFP) under the regulation of the *dei*^*ChO-1353*^ enhancer, while the attachment cells expressed red fluorescent protein (RFP) under the regulation of the *dei*^*attachment*^ enhancer (Halachmi et al., 2016). One of the 31 genes identified in the screen as being required for normal morphogenesis of the larval LCh5 organ was *shaven* (*sv*), which encodes for the *Drosophila* Pax2 homologue D-Pax2. The knockdown of *sv* within the ChO lineage led to loss of expression of the *dei*^*ChO-1353*^ reporter from the cap cells, suggesting that D-Pax2 is a positive regulator of *dei* transcription in this cell type (Hassan et al., 2018).

The second screen was a yeast one hybrid (Y1H) screen aimed at identifying proteins that bind *directly* to the *dei*^ChO-389^ enhancer. The screen was performed on a *D. melanogaster* whole embryo library using the *dei*^ChO-389^ sequence as a bait. Two proteins were identified to bind the bait with very high confidence in the interaction: D-Pax2 and LamC. Three additional candidates were identified as binders with moderate confidence in the interaction: Fax, Lola and Toy (Figure S1). Together, the results of the two independent screens identified D-Pax2 as a putative direct transcriptional activator of *dei* and an important player in ChO development.

### D-Pax2/Sv activates *dei* expression in the cap cell

To validate the *sv* RNAi-induced phenotype, we characterized the LCh5 organs of *sv* mutant embryos (*sv*^*6*^, *sv*^*7*^). In accordance with the knockdown phenotypes, the expression of *dei* in the cap cell was abolished in *sv-*deficient embryos (Figure 2A-B). The expression of additional genes associated with proper differentiation of the cap cell, such as *αTub85E*, was reduced as well (Figure 2C-D). The loss of *sv* did not eliminate the expression of either scolopale-specific proteins (Crumbs, Pros) or neuronal markers (Futsch, Elav, Nrg), suggesting that it did not affect primary cell fate decisions, however its loss prevented normal morphogenesis of the sensory unit (Figure 2E-H).

**Figure 2.**
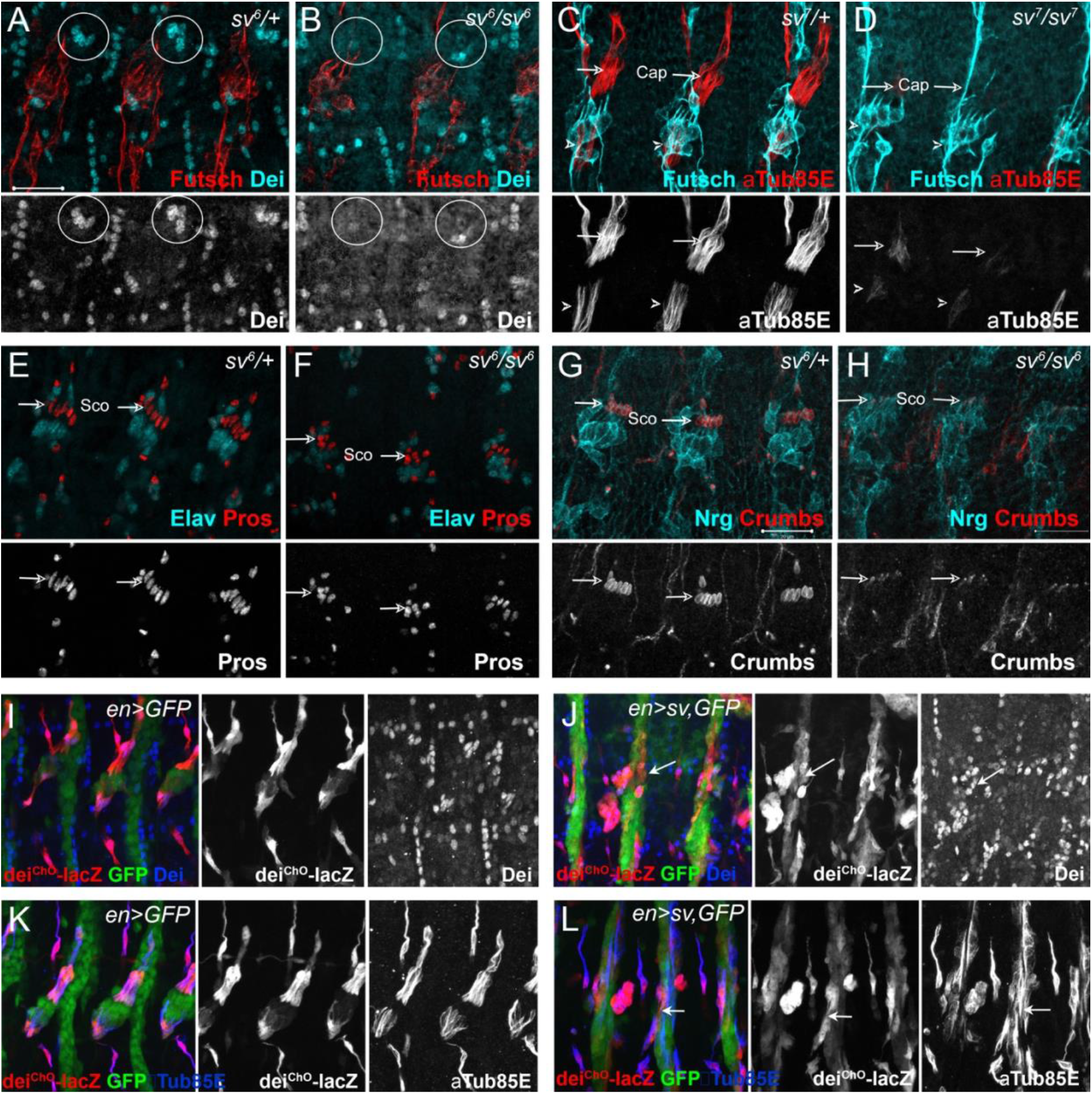
Sv/D-Pax2 activates *dei* expression in the cap cell. (A-B) Representative abdominal segments of stage 16 embryos stained for Dei (cyan) and the neuronal marker Futcsh (red). The anti-Dei staining is shown separately in the lower panel. (A) A *sv*^*6*^ heterozygous embryo demonstrating the expression of Dei in the cap cell nuclei (circled). (B) The expression of Dei is lost in homozygous *sv*^*6*^ embryos. (C-D) Heterozygous (C) and homozygous (D) *sv*^*7*^ embryos stained for αTub85E (red) and Futsch (cyan). The arrows mark the cap cells; the arrowheads mark the ligament cells. The anti-αTub85E staining is shown separately in the lower panel. (E-F) Heterozygous (E) and homozygous (F) *sv*^*6*^ embryos stained for Elav (cyan) and Pros (red). The arrows mark the nuclei of scolopale cells. The anti-Pros staining is shown separately in the lower panel. (G-H) Heterozygous (G) and homozygous (G) *sv*^*6*^ embryos stained for Nrg (cyan) and Crumbs (red). The arrows mark the nuclei of scolopale (Sco) cells. The anti-Crumbs staining is shown separately in the lower panel. (I-L) Representative abdominal segments of stage 16 embryos that express GFP (I, K) or GFP and *sv* (J, L) under the regulation of *en-Gal4*. The embryos carry the *dei*^*ChO-262*^*-lacZ* marker (anti βGal staining is shown in red) and are stained with anti-Dei (blue in I-J) or anti-αTub85E (blue in K-L). The staining patterns of *dei*^*ChO-262*^*-lacZ*, Dei and αTub85E are shown separately on the right.

To further test the ability of D-Pax2/Sv to activate *dei* expression *in vivo*, we ectopically expressed *sv* under the regulation of *en-Gal4* and examined the resulting changes in gene expression patterns. The ectopic expression of *sv* led to ectopic expression of both the endogenous *dei* gene and the *dei*^*ChO-262*^ transcriptional reporter, as well as the cap cell marker αTub85E (Figure 2I-L). The morphology of the *sv*-overexpressing LCh5 organs was abnormal. These observations corroborate the notion that D-Pax2/Sv is an activator of *dei* that plays a critical role in ChO morphogenesis.

### Pros represses *dei* in the scolopale cell

The *sv* gene is expressed broadly within the ChO lineage during early stages of organ development and is then gradually restricted to the cap and scolopale cells (Czerny et al., 1997; Fu et al., 1998), where its expression level remains high during late embryogenesis and larval stages. Yet, the expression of its downstream target gene *dei* is, normally, activated in the cap cell but is excluded from the scolopale cell. This discrepancy in the expression pattern of D-Pax2/Sv and *dei* may point to the presence of a scolopale-specific repressor that prevents the transcription of *dei* in this D-Pax2/Sv-expressing cell. The results of the abovementioned RNAi screen identified the transcription factor Pros as a good candidate for being that repressor. *pros* and *sv* RNAi had opposite effects on the expression of the *dei*^*ChO-1353*^-*GFP* reporter. Whereas the knockdown of *sv* led to a loss of the reporter expression from the cap cell, the knockdown of *pros* led to expansion of its expression into the scolopale cell suggesting that, normally, Pros represses *dei* in this cell (Hassan et al., 2018).

To validate the *pros* knockdown phenotypes and to further test the idea that Pros acts as a repressor of *dei* in the scolopale cell, we characterized the ChOs of *pros*^*17*^ embryos. As seen in Figure 3A-B, the loss of *pros* led to ectopic expression of both the endogenous *dei* gene and the *dei*^*ChO-262*^ reporter, as well as the cap cell marker αTub85E in the scolopale cells. The *pros*-deficient scolopale cells developed into cap-like cells rather than ligament-like or attachment cell-like cells. This was indicated by the upregulation of *dei* which was not accompanied by upregulation of the transcription factor Sr that is normally co-expressed with Dei in the ligament and attachment cells but is excluded from the cap cells (Figure 3C-D). The observed alterations in gene expression pattern do not reflect a full scolopale-to-cap cell fate transformation, as the Pros-deficient scolopale cells still maintain some of their scolopale-specific characteristics, such as Eyes Shut and α–Catenin expression (Figure 3E-H). As the number of neurons, ligament, cap, and cap-attachment cells remained normal in *pros* mutant embryos (Figure S2), we conclude that Pros does not influence primary cell-fate decisions within the LCh5 lineage.

**Figure 3.**
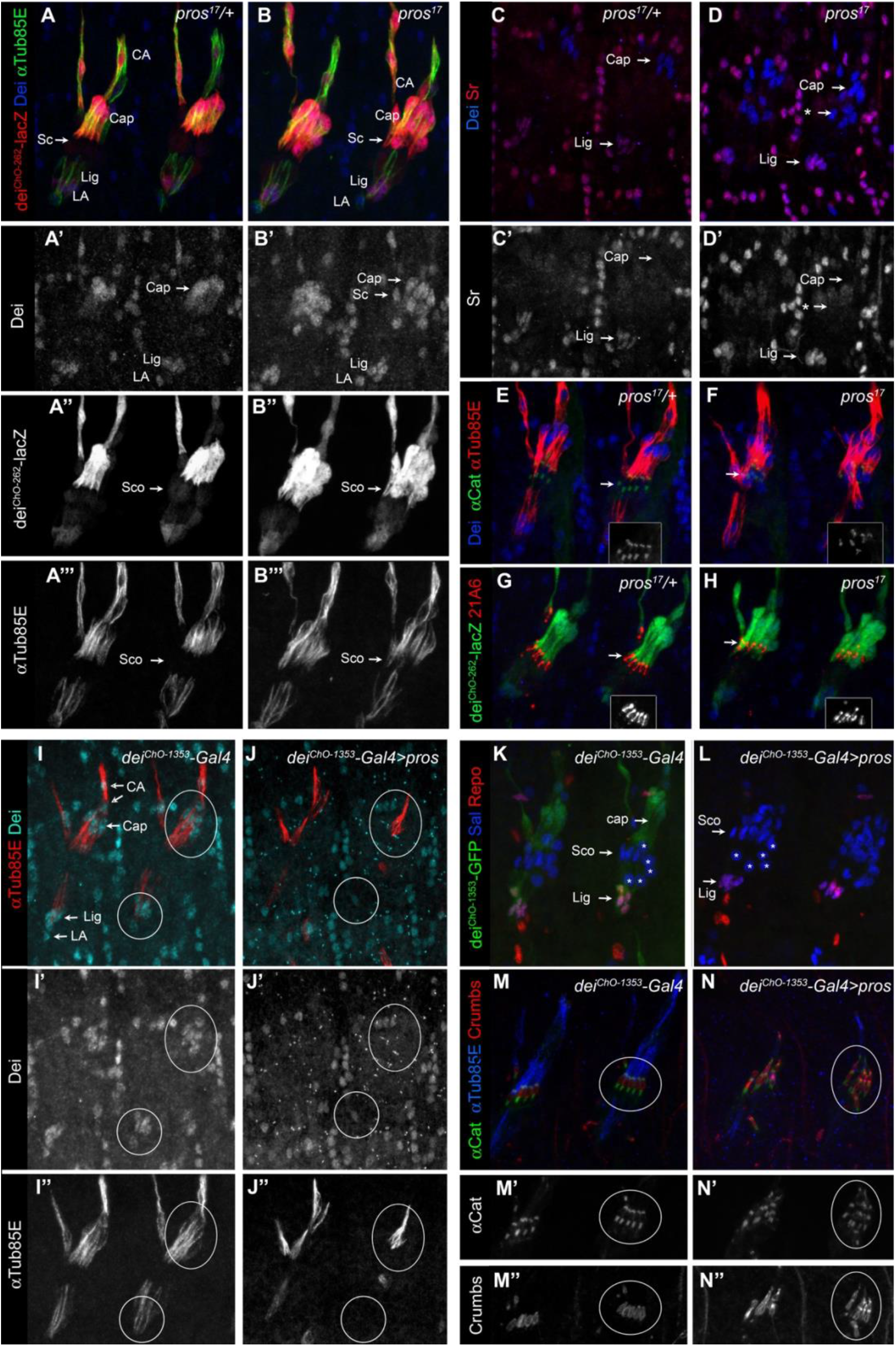
Pros represses *dei* in the scolopale cell. (A-B) Representative abdominal segments of stage 16 *pros*^*17*^ heterozygous (A) and homozygous (A) embryos carrying the *dei*^*ChO-262*^*-lacZ* marker (shown in red) and stained for Dei (blue) and αTub85E (green). Each of the three channels is shown separately below. Note the expansion of Dei and the *dei*^*ChO-262*^*-lacZ* marker into the scolopale cells in *pros*^*17*^ mutant embryos. The arrows point to the scolopale cells. (C-D) *pros17* heterozygous (C) and homozygous (D) embryos stained for Dei (blue) and Sr (red). Note that the ectopic expression of Dei in the scolopale cells of pros mutant embryo is not accompanied by ectopic expression of Sr (the arrow labeled with asterisks in D). (E-F) *pros*^*17*^ heterozygous (E) and homozygous (F) embryos carrying an α-Catenin-GFP reporter and stained for Dei (blue, shown separately in the inset) and αTub85E (red). (G-H) *pros*^*17*^ heterozygous (G) and homozygous (H) embryos carrying the *dei*^*ChO-262*^*-lacZ* reporter (shown in green) and stained with the scolopale marker anti-Eyes Shut (MAb21A6, red, shown separately in the inset). Note that the expression of both Eyes Shut/21A6 and α-Catenin is maintained in the *pros*-deficient scolopale cells. (I-J) *dei*^*ChO-1353*^*-Gal4* (I) and *dei*^*ChO-1353*^>*UAS-pros* (J) embryos stained for Dei (cyan, shown separately in I’-J’) and αTub85E (red, shown separately in I’’-J’’). The cap and ligament cells are circled. Note the loss of Dei and αTub85E expression upon Pros expression. (K-L) *dei*^*ChO-1353*^ (I) and *dei*^*ChO-1353*^>*UAS-pros* (J) embryos carrying the *dei*^*ChO-1353*^*-lacZ* reporter (shown in green) stained for Sal (blue) and Repo (red). (M-N) *dei-Gal4* (M) and *dei-Gal4>UAS-pros* (N) embryos carrying an α-Catenin-GFP reporter (green, shown separately in M’-N’) and stained for Crumbs (red, shown separately in M’’-N’’) and αTub85E (blue). Note the duplication of scolopale-specific structures in the cap cell expressing Pros ectopically (circled).

The ability of Pros to repress *dei* was not restricted to the scolopale cell. Ectopic expression of *pros* in the LCh5 lineage, using a *dei*^*ChO-1353*^*-Gal4* driver, abolished the expression of both the endogenous *dei* gene and a *dei*^*ChO-1353*^*-GFP* reporter in the cap and ligament cells as well (Figure 3I-L). In parallel to the repression of *dei*, ectopic expression of *pros* was sufficient for upregulating scolopale-specific genes and, moreover, for driving the formation of ectopic scolopale-specific structures within the affected cap cells. Most notably, the Pros-expressing cap cells manifested scolopale rods and ectopically expressed Crumbs and α-Catenin in a scolopale-characteristic pattern (Figure 3M-N).

### D-Pax2/Sv and Pros regulate the transcription of *dei* via *dei*^*ChO-262*^

The fact that the expression of the *dei*^*ChO*^ transcriptional reporters was affected similarly to the endogenous *dei* gene by both *sv* and *pros* loss- and gain-of-function, suggested that both D-Pax2/Sv and Pros regulate *dei*’s transcription in the ChOs through this regulatory module. To test this hypothesis, we deleted the *dei*^*ChO-262*^ region from the fly genome by CRISPR/Cas9-mediated genome editing, resulting in a new regulatory allele of *dei* (*dei*^Δ*ChO*^). As expected, in homozygous *dei*^Δ*ChO*^ embryos the expression of *dei* was lost from the cap and ligament cells but remained intact in the rest of the *dei*-expressing cells: CA, LA and tendon cells (Figure 4A-B). This observation strongly suggests that the *dei*^*ChO-262*^ enhancer constitutes the sole regulatory module driving *dei* expression in the cap and ligament cells.

**Figure 4.**
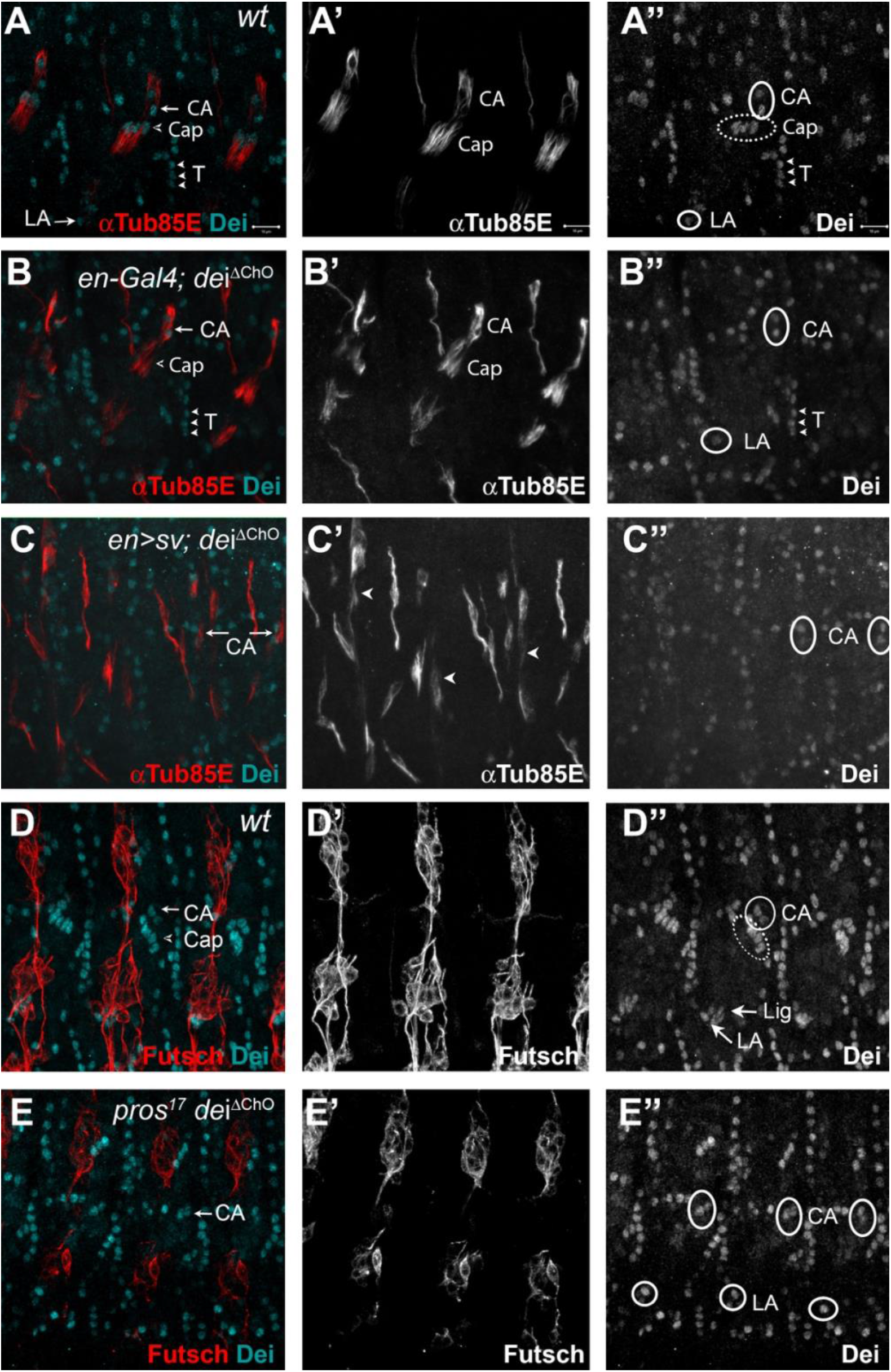
D-Pax2/Sv and Pros regulate the transcription of *dei* via the *dei*^*ChO-262*^ regulatory module. (A-B) Stage 16 *wt* (A) and *dei*^*ΔChO*^ (B) embryos stained for Dei (cyan) and αTub85E (red). Note that in (B) Dei is still evident in the cap-attachment and ligament-attachment cells (CA and LA, arrows in A, circled in A’’), as well as in tendon cells (arrowheads), but is lost from the cap and ligament cells. (C) An embryo in which *sv* was expressed under the regulation of *en-Gal4* in a *dei*^*ΔChO*^ background. Sv was unable to induce Dei expression in the cap and ligament cells in the absence of the *dei*^*ChO-262*^ enhancer. Ectopic expression of αTub85E is evident within the *en* domain (arrowheads in C’). (D-E) *wt* (D) and A *pros*^*17*^ *dei*^*ΔChO*^ homozygous embryo (E) stained for Dei (cyan) and anti-Futsch (red). The *pros*^*17*^ *dei*^*ΔChO*^ embryo (E) presents a *dei*^*ΔChO*^-like and not *pros*^*17*^-like expression pattern of the Dei protein, indicating that the *dei*^*ΔChO*^ deletion is epistatic to *pros* loss-of-function. The neurons present the typical *pros* axonal pathfinding defects.

To further test the notion that D-Pax2/Sv regulates *dei* expression via the *dei*^*ChO-262*^ enhancer we examined the ability of D-Pax2/Sv to activate *dei* expression in the *dei*^Δ*ChO*^ background. We found that ectopic expression of D-Pax2/Sv failed to induce ectopic *dei* expression in the *dei*^Δ*ChO*^ background (Compare Figure 4C and 2L), indicating that the *dei*^*ChO-262*^ enhancer is indispensable for the ability of D-Pax2/Sv to activate *dei* transcription. In a complementary experiment, we tested the effect of *pros* loss-of-function on the expression of *dei* in the *dei*^Δ*ChO*^ background. We found that the *dei*^*ΔChO*^ regulatory mutation was epistatic to *pros* loss-of-function, so that no ectopic expression of *dei* was observed in the *pros*-deficient scolopale cells in embryos homozygous for the *dei*^Δ*ChO*^ mutation (Compare Figure 4D-E and Figures 3B, D). Based on these results we concluded that the expression of *dei* in ChOs depends on the presence of the *dei*^*ChO-262*^ regulatory region and that this enhancer integrates the positive and negative inputs of D-Pax2/Sv and Pros, respectively, to drive a lineage-specific *dei* expression.

### D-Pax2/Sv and Pros are direct transcriptional regulators of *dei*

Our genetic analyses revealed that D-Pax2/Sv and Pros regulate the expression of *dei* in opposing manners through the function of the *dei*^*ChO-262*^ enhancer. In addition, the Y1H screen identified a direct interaction between D-Pax2/Sv and *dei*^*ChO-389*^ (which include the *dei*^*ChO-262*^ module). We therefore hypothesized that the *dei*^*ChO-262*^ enhancer contains binding sites for D-Pax2/Sv and Pros. Motif search analysis predicted that the *dei*^*ChO-262*^ sequence encodes one canonical binding site for D-Pax2/Sv (Figure 5B, D-Pax2/Sv Site 2). On the other hand, we were unable to predict binding sites for Pros in the *dei*^*ChO-262*^ sequence.

**Figure 5:**
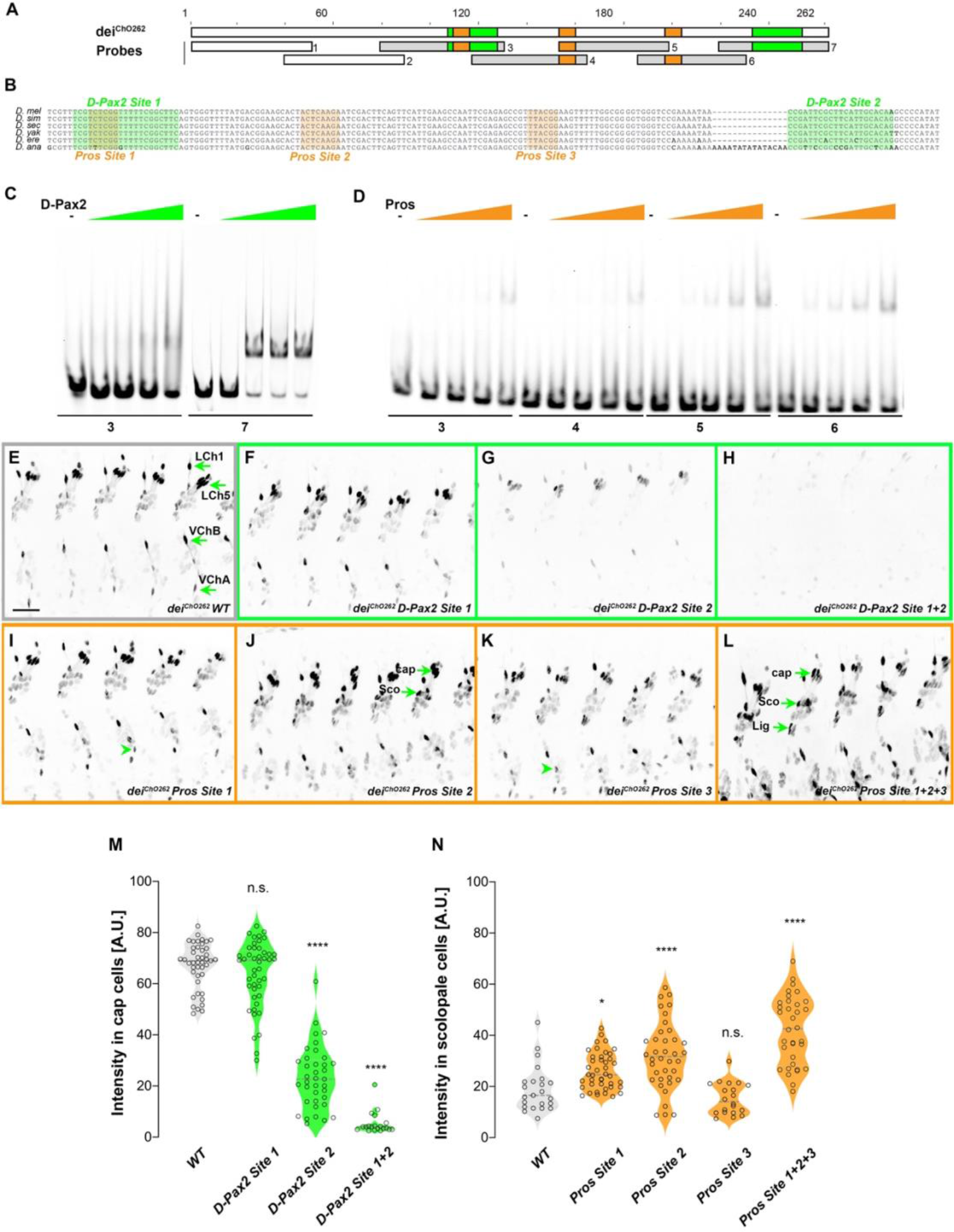
*dei*^*ChO-262*^ contains two binding sites for the activator D-Pax2/Sv and three binding sites for the repressor Pros. (A) Schematic representation of the *dei*^*ChO-262*^ enhancer. Green and orange boxes represent the location of the D-Pax2/Sv and Pros binding sites identified by systematic EMSAs using oligos corresponding to the regions represented by boxes 1-7. (B) Sequence alignment between six Drosophila species for the region of the *dei*^*ChO-262*^ enhancer containing the three D-Pax2/Sv and Pros sites (labeled and highlighted in green and orange, respectively). Dashes indicate gaps in the aligned sequence. (C) D-Pax2/Sv binds to two sites in *dei*^*ChO-262*^, one low affinity site in fragment 3 (*D-Pax2 site 1*) and one high affinity site in fragment 7 (*D-Pax2 site 2*), as demonstrated with EMSA. The full screen is shown in Supplementary Figure S3. (D) Pros binds to three sites in *dei*^*ChO-262*^: *Pros site 1* in fragment 3, *Pros site 2* in an overlapping sequence in fragments 4 and 5, *Pros site 3* in fragment 6, as demonstrated with EMSA. The full screen is shown in Supplementary Figure S4. (E-L) Expression of wild-type (E) and mutated (F-L) *dei*^*ChO-262*^*-lacZ* reporter constructs in abdominal segments A2-A6 of representative stage 16 embryos. The name of the construct is indicated in the bottom of each panel. The green arrows in E point to the cap cells of the various ChO of one abdominal segment: LCh1-lateral ChO1, LCh5-pentascolopidial organ, VChB and VChA are two ventral ChOs. The green arrows in J and L point to cap, scolopale (Sco) and ligament (Lig) cells of LCh5 organs where elevated level of reporter expression is evident. The arrowheads in I and K point to ligament cells of the ventral ChOs. in (M-N) Quantification of reporter activity in nuclei of cap (M) and scolopale cells (N) from LCh5 of segment A2 in embryos carrying the indicated constructs (n=10 embryos for each genotype). In violin plots, each point represents an individual nucleus, median is shown as dark gray dashed line. Asterisks denote significant difference from wild-type activities, (*) - P<0.05, (****) – P<0.0001, n.s. – not significant (Dunnett’s multiple comparison test).

To test whether D-Pax2/Sv actually binds to the predicted site, and to search for additional D-Pax2/Sv binding sites and for Pros binding sites, we systematically screened the *dei*^*ChO-262*^ sequence with electrophoretic mobility shift assays (EMSAs, Figure 5 and Supplementary Figures S3 and S4). As shown in Figure 5C, purified D-Pax2/Sv DNA-binding domain bound strongly to the region containing the predicted binding site (probe 7). In addition, we identified another region (in probe 3), that bound D-Pax-2/Sv at a lower affinity (Figure 5C and Supplementary Figure S3). Using purified Pros-S DNA-binding domain we found that four fragments - probes 3, 4, 5 and 6 - bound Pros *in vitro* (Figure 5D). Comprehensive mutagenesis and competition assays revealed three Pros conserved binding sites within these fragments (Supplementary Figure S4), one of them partially overlapping the low-affinity binding site of D-Pax2/Sv (Figure 5B). Interestingly, none of these identified binding sites resemble the known binding sites for Pros identified by SELEX (Hassan et al., 1997), by functional studies (Cook et al., 2003), or by single-cell omics analyses (Bravo González‐Blas et al., 2020).

To test the *in vivo* role of the D-Pax2/Sv sites identified *in vitro,* we mutated them either individually, or in combinations, in the context of a *dei*^*ChO-262*^ reporter transgene encoding for nuclear β-galactosidase. Mutation of the canonical D-Pax2/Sv site 2 reduced the expression level driven by *dei*^*ChO-262*^ in cap cells (Figure 5G and 5M). Mutation of D-Pax2/Sv site 1 had no significant effect on its own (Figure 5F and 5M) but led to a complete suppression of *dei*^*ChO-262*^ function when combined with a mutation in D-Pax2/Sv site 2 (Figure 5F and 5M). These results suggest that D-Pax2/Sv regulates the expression of *dei* in cap cells by binding to two D-Pax2/Sv binding sites within the *dei*^*ChO-262*^ enhancer.

We next tested whether the Pros sites identified by EMSA function *in vivo* to suppress *dei*^*ChO-262*^ activity in scolopale cells. Mutating the Pros site 2 resulted in ectopic expression of the reporter gene in the scolopale cells of LCh5 (Figure 5J and 5M). In contrast, disruption of either Pros site 1 or site 3 had no detectable effects on the expression of *dei*^*ChO-262*^ in the scolopale cells of LCh5 (Figure 5I, 5K and 5M) but induced ectopic expression in scolopale cells of the LCh1 and VChA/B organs (Figure 5I and 5K, arrowheads). In addition, the disruption of Pros site 3 led to elevation in the reporter’s level in the LCh5 ligament cells (Figure 5K). Simultaneous mutation of all three Pros binding sites in *dei*^*ChO-262*^ resulted in intensified ectopic expression in the scolopale cells of all ChOs (Figure 5L and 5N). Based on these results we conclude that Pros represses *dei* expression in the scolopale cell by binding to three low-affinity binding sites in *dei*^*ChO-262*^ that function additively.

### The *dei*^*ChO-262*^ enhancer is essential for normal ChO function and larval locomotion

We have previously shown that the cap cell plays a crucial role in propagating muscle-generated mechanicals signals to the sensory neuron (Hassan et al., 2019). To test whether *dei* expression in the cap and ligament cells, mediated solely by the *dei*^*ChO-262*^ enhancer, is essential for the proprioceptive function of the ChO, we analyzed the pattern of locomotion of freely moving *dei*^Δ*ChO*^ larvae and compared it to wild-type larvae and larvae homozygous for the *dei*^*KO-mCherry*^ null allele. As shown in Figure 6, *wild-type* larvae crawled persistently 94.5 ± 12.4% (n=27) of the time with a very few changes of direction or head swipes (Figure 6A, E, H, and Video S1-2). In contrast, the *dei*^*KO-mCherry*^ larvae exhibited frequent changes of moving direction and longer pauses, walking on average only 38.9 ± 19.4% (n=25) of the time. While pausing, the *dei*^*KO-mCherry*^ larvae swiped their heads extensively (Figure 6B, F, H, and Video S3-4). The *dei*^Δ*ChO*^ larvae exhibited locomotion phenotypes that were similar to, though slightly less severe and more variable than, those of the *dei*^*KO-mCherry*^ larvae (Figure 6C, G and H, Video S5-6). On average, the *dei*^Δ*ChO*^ larvae crawled 51.9 ± 24.9% (n=24) of the time and swiped their heads often; in 15.8% of the time their body angle was higher than 250° or lower than 110°, compared to 9.9% in *dei*^*KO-mCherry*^ and 1.2% of the time in *wt* larvae (Figure 6H). These results demonstrate that the *dei*^*ChO-262*^ regulatory element is crucial for proper function of the ChO as its deletion resulted in sensory dysfunction and uncoordinated movement.

**Figure 6.**
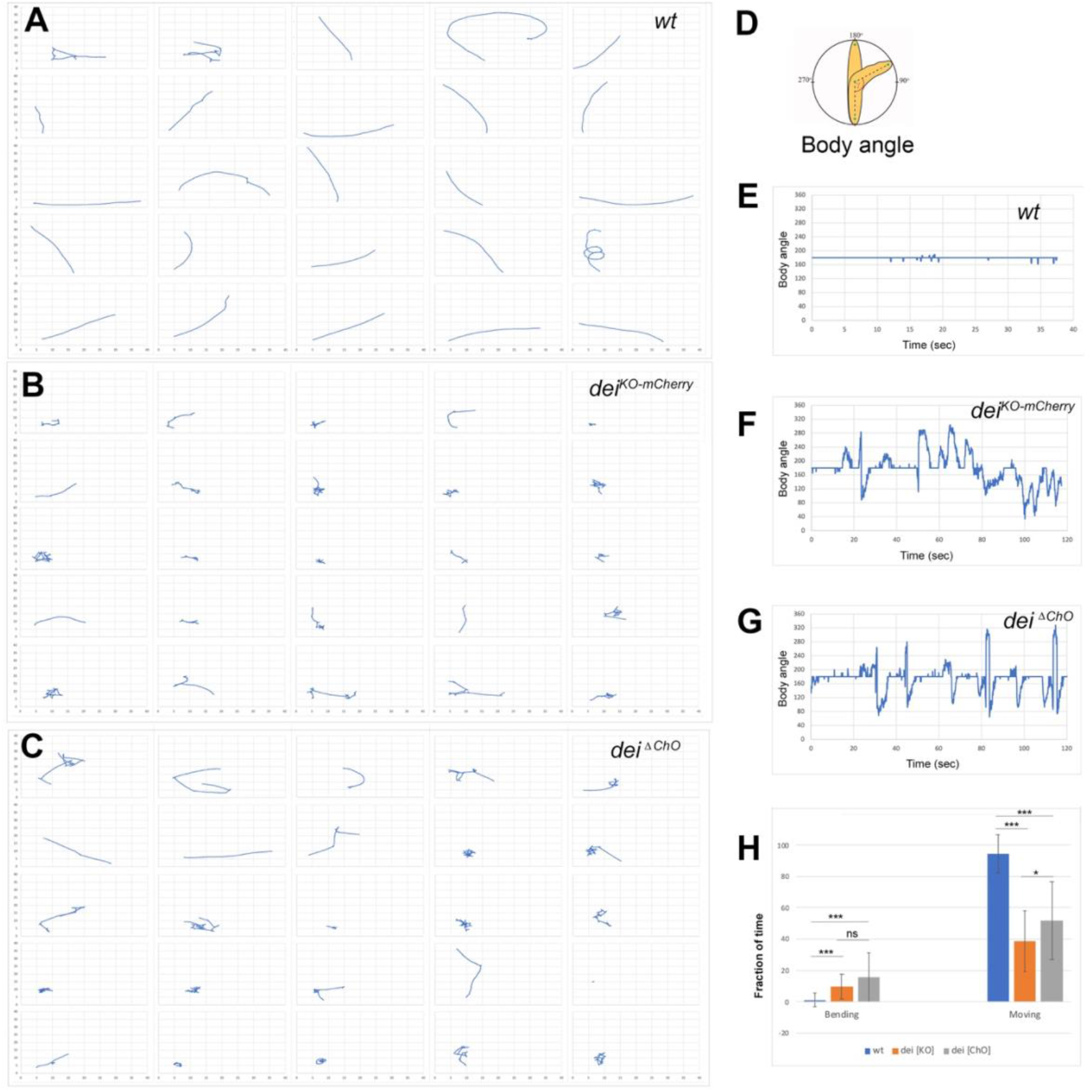
The *dei*^*ChO-262*^ enhancer is essential for normal larval locomotion. (A-C) Crawling trajectories of 25 wild-type (A) *dei*^*KO-mCherry*^ (B) *and dei*^*ΔChO*^ (C) larvae. Each trajectory is shown in a square that represents 40×40mm area. (D) schematic representation of the body angle, γ, defined as the angle between the head and the body axis. (E-G) Representative time evolutions of the body angle of a wild-type (E), *dei*^*KO-mCherry*^ (F) and *dei*^*ΔChO*^ (G) larvae. The wild-type larva walks persistently, and the body angle stays 180 degrees most of the time (a 40 sec interval is shown, after which the larva exited the filmed arena). The *dei*^*KO-mCherry*^ and *dei*^*ΔChO*^ mutant larvae display frequent changes in the direction of motion and long pauses accompanied by extensive head swiping (120 sec intervals are shown). (H) A graph showing the average fraction of the time the larvae were crawling (GoPhase) and the fraction of time in which the body was bended more than 70 degrees (the measured angle γ was higher than 250 degrees or lower than 110 degrees). n=24-27 for all genotypes; error bars represent the standard deviation. *** p<0.0001, * p<0.05, ns= non-significant. P values were calculated using the unpaired two-tailed Mann-Whitney test.

## DISCUSSION

### Opposing activities of D-Pax2 and Pros dictate cap versus scolopale differentiation programs by regulating the *dei* gene

In this work we identify a small GRN that governs the alternative differentiation programs of two ‘cousins once removed’ cells within the ChO lineage - the cap cell and the scolopale cell. We show that Pros and D-Pax2/Sv are direct regulators of *dei* that together dictate its expression in the cap cell and its repression in the scolopale cell. Both D-Pax2/Sv and Pros exert their effects on *dei* transcription via a 262 bp chordotonal-specific enhancer (*dei*^*ChO-262*^) in which two D-Pax2/Sv and three Pros binding sites were identified. Following primary cell fate decisions within the ChO lineage, Pros expression becomes restricted to the scolopale cell (Doe et al., 1991; Vaessin et al., 1991), whereas D-Pax2/Sv expression becomes restricted to the scolopale and cap cells (Figure 1A), similar to its behavior in the external sensory lineages (Johnson et al., 2011; Kavaler et al., 1999). D-Pax2/Sv activates the expression of *dei* in the cap cell but is unable to do so in the scolopale cell where Pros is co-expressed. If D-Pax2/Sv activity is compromised, the cap cell fails to express *dei* and loses some of its differentiation markers, such as the expression of αTub85E. In contrast, if Pros activity is lost, *dei* is ectopically expressed in the scolopale cell that, as a consequence, acquires some cap cell features including the expression of αTub85E (Figure 7). The observed D-Pax2/Sv- and Pros-associated phenotypes do not reflect genuine cell fate transformations, suggesting that D-Pax2/Sv and Pros do not affect primary cell fate decisions within the ChO lineage. Rather, the observed phenotypes reflect a failure of the cap and scolopale cells to follow the cell type-specific differentiation programs responsible for bringing about their characteristic cellular phenotypes. In the adult external sensory lineage Pros was shown to be important for the specification of the pIIb precursor, which gives rise to the neuron and sheath cell (the scolopale counterpart). However, the absence of Pros from the pIIa precursor, which gives rise to the hair and socket cells (the cap and cap-attachment cells counterparts) was even more critical for proper development of this branch of the lineage (Manning and Doe, 1999). This phenomenon is somewhat conserved in the larval ChO. While Pros is required for proper differentiation of the scolopale cell, its absence from the cap cell is critical for adopting the correct differentiation programs within the lineage.

**Figure 7.**
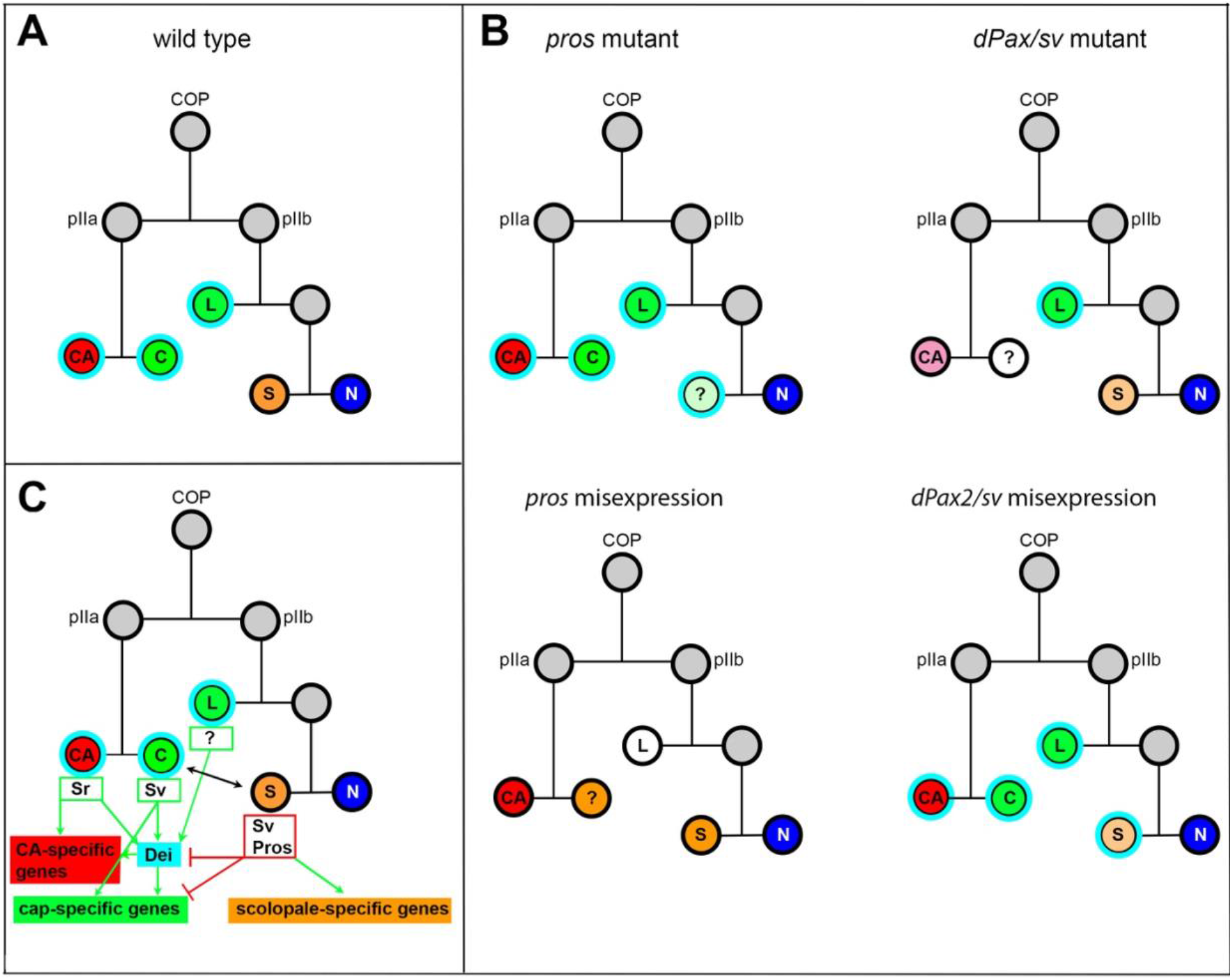
Summary of the relations between Sv, Pros and Dei in the ChO lineage and their effect on ChO development. (A) A *wt* ChO lineage. The cap (C) and ligament (L) cells are depicted in green, the scolopale cell (S) is depicted in orange and the neuron (N) is depicted in blue. Cells that expressed Dei are circled in light blue. (B) the loss of pros leads to upregulation of Dei in the scolopale cell and to failure of scolopale cell differentiation. In contrast, the loss of Sv leads to loss of Dei expression from the cap cell and failure of cap cell differentiation. The Sv-deficient scolopale cells are also abnormal. The CA cells which depend on the cap cell for their development/maintenence also appear abnormal is *sv* mutants. Misexpression of Pros leads to repression of Dei in the cap and ligament cells, preventing their normal differentiation. The Pros-expressing cap cells adopt some scolopale-specific features. In contrast, over-expression of Sv leads to ectopic expression of Dei. Due to the presence of Pros, the level of expression of *dei* in the scolopale cell is restricted. (C) A schematic summary showing the relations between Pros, Sv and Dei and their relations to cell type-specific differentiation programs. In the CA cells, *dei* is activated by Sr via the *dei*^*attachment*^ enhancer. Both Sr and Dei are required there for the activation of CA-specific genes. In the cap cell *dei* is activated by Sv via the *dei*^*ChO-262*^ enhancer. Sv is required for activating cap-specific genes in both Dei-dependent and independent ways. In the scolopale cells *dei* is repressed by Pros via the *deiChO-262* enhancer. Pros is required in addition for activating scolopale-specific genes. Dei is also expressed in the ligament cells and is required for their correct differentiation. The regulators of *dei* in the ligament cell are yet to be identified.

Opposing effects of D-Pax2 and Pros activities on cell differentiation have been previously identified in the regulation of neuronal versus non-neuronal cell fate decisions in the developing eye, where they play a role in modulating the Notch and Ras signaling pathway (Charlton-Perkins et al., 2011). Interestingly, in the R7 equivalence group Pros and D-Pax2 can only alter the cell type-specific differentiation program of cells that already express the other gene (Charlton-Perkins et al., 2011). Similarly, in the ChO lineage, ectopic expression of Pros in the cap and ligament cells transforms the D-Pax2-positive cap cell towards a scolopale cell identity but does not affect the D-Pax2-negative ligament cell in a similar fashion, even though the ectopic expression of Pros does repress the transcription of *dei* in both cell types. Additionally, a loss of Pros activity in the scolopale cell can transform the identity of this cell towards a cap cell identity only in the presence of D-Pax2.

We do not know how Pros opposes the effect of D-Pax2 and inhibits the expression of *dei* in the scolopale cell. The results of the DNA-binding assays suggest that the inhibitory effect of Pros is not mediated through binding competition with D-Pax2 at the D-Pax2 high-affinity site (site 2), since this site does not overlap with a Pros-binding site (Figure 5B). The D-Pax2 low-affinity site does overlap with a Pros binding site, however, while being important for robust *dei* expression, this site is dispensable in the presence of the high-affinity site. It is possible that binding of Pros to the *dei*^*ChO-262*^ enhancer targets this sequence to a repressed heterochromatin domain as was recently shown for other Pros target genes in differentiating neurons (Liu et al., 2020).

### How is *dei* regulated in other ChO cell types?

*dei* is expressed in four out of six cell types comprising the ChO: the cap-attachment and ligament-attachment cells, in which *dei* transcription is activated by Sr via the *dei*^*attachment*^ regulatory module (Nachman et al., 2015), and the cap and ligament cells in which the expression of *dei* is regulated via the *dei*^*ChO-262*^ enhancer. We now show that D-Pax2 activates *dei* transcription in the cap cell, and that Pros inhibits its expression in the scolopale cell. The identity of the positive regulator/s of *dei* in the ligament cell, whose cell-fate is determined by the glial identity genes *gcm* and *repo* (Campbell et al., 1994; Halter et al., 1995; Jones et al., 1995), and the identity of the negative regulator/s of *dei* in the neuron remains unknown. Interestingly, the expression of *dei* was found to be altered in response to ectopic expression of *gcm* in the embryonic nervous system; its expression was upregulated at embryonic stage 11, but was repressed in embryonic stages 15-16 (Egger et al., 2002). This observation points to GCM as a potential regulator of *dei* expression in the ligament cells. Another interesting candidate for repressing *dei* in the sensory neuron is the transcriptional repressor Lola. Lola has been identified as a putative direct regulator of *dei* in the Y1H screen and was shown to be required in post-mitotic neurons in the brain for preserving a fully differentiated state of the cells (Southall et al., 2014). The possible involvement of Gcm and Lola in the regulation of *dei* awaits further studies. The observed upregulation of the *dei*^*ChO-262*^ reporter in the ligament cells of embryos with mutated Pros binding sites may reflect an early role of Pros in the pIIb precursor before its restriction to the scolopale cell, which prevents *dei* expression in the ligament cell.

### The *dei*^*ChO-262*^-driven Dei expression is critical for organ functionality

Although the loss of *dei* in the genetic/cellular milieu of the ligament cell (unlike the cap cell), even when accompanied by ectopic expression of Pros, is not sufficient for transforming ligament cell properties towards those of scolopale cells, we know that the expression of *dei* in the ligament cell is critical for its proper development. Ligament-specific knockdown of *dei* leads to failure of the ligament cells to acquire the right mechanical properties and leads to their dramatic over-elongation (Hassan et al., 2018). By analysing the locomotion phenotypes of larvae homozygous for a *dei* null allele and the newly generated cap and ligament-specific *dei*^Δ*ChO*^ allele, we could show that the expression of *dei* in the cap and ligament cells is crucial for normal locomotion. Thus, we conclude that the correct expression of *dei* within the ChO is critical for organ functionality. Surprisingly, the gross morphology of LCh5 of *dei*^Δ*ChO*^ larvae appears normal (Data not shown). Yet, in a way that remains to be elucidated, the Dei-deficient cap and ligament cells fail to correctly transmit the cuticle deformations to the sensory neuron, most likely due to changes in their mechanical properties.

## MATERIALS AND METHODS

### Fly strains

The following mutant and reporter alleles of *sv* and *pros* were used: *sv*^*6*^/*act-GFP*, *sv*^*7*^/*act-GFP*, (Kavaler et al., 1999), *Dpax2*^*D1*^*-GFP* (Johnson et al., 2011), *pros*^*17*^/*TM6B, Tb*^*1*^ (BDSC:5458). The following Gal4 drivers and UAS strains were used: *dei*^*ChO-1353*^*-GFP,dei*^*attachment*^*-RFP;en-Gal4* (Halachmi et al., 2016), *en-Gal4* (Brand and Perrimon, 1993), *P{UAS-3xFLAG-pros.S}14c, y1 w*; Pin^1^/CyO* (BDSC:32245). *UAS-sv-RNAi* (VDRC:107343), *UAS-sv* (Kavaler et al., 1999), UAS-D-α-Catenin-GFP (Oda and Tsukita, 1999).

The *dei*^*ChO*^*-Gal4* driver was constructed by cloning the *dei*^*ChO-1353*^ regulatory module described in (Nachman et al., 2015) into the *pChs-gal4* vector which was then used for the generation of transgenic fly strains (insertions are available on the X, 2^nd^ and 3^rd^ chromosome). The *dei*^*ChO-262*^-*lacZ* strain was generated as previously described in (Nachman et al., 2015). For mutational analysis of the putative binding sites, wild-type and mutated *dei*^*ChO-262*^ fragments were synthesized by GenScript (USA) and cloned into the reporter constructs *placZattB* (Table S4). Plasmids were integrated into the *attP2* landing site by BestGene Inc (Chino Hills CA, USA).

The *dei*^*ΔChO*^ allele was generated by GenetiVision (Houston TX, USA) via multiplex targeting with two sgRNAs: 5’GCACTTGTTTGCGTTTACATTAC3’ and 5’GGCGAGAAGTATTCCCTGCG3’; creating a defined deletion of 307 bps spanning the *dei*^*ChO-262*^ fragment. The presence of the desired deletion was verified by sequencing.

### Embryo staining and image analysis

Immuno-staining of whole-mount embryos was performed using standard techniques. The following primary antibodies used in this study were: Rabbit anti Sv/D-Pax2 (1:10,000) (Johnson et al., 2011), Rabbit anti α85E-Tubulin (1:200) (Klein et al., 2010), Mouse anti α85E-Tubulin (1:20) (Nachman et al., 2015), Rabbit anti Dei (1:50)(Egoz-Matia et al., 2011), Rabbit anti Spalt (1:500) (Halachmi et al., 2007), Rat anti NRG (1:1000) (Banerjee et al., 2006), and Mouse anti-βGal (1:1000, Promega). The following antibodies were obtained from the Developmental Studies Hybridoma Bank: Mouse anti Pros (MR1A, 1:20), Mouse anti Futsch (22C10, 1:20), Rat anti-ELAV (7E8A10, 1:50), Mouse anti-Eys (21A6, 1:20), Mouse anti-Repo (8D12, 1:10), Mouse anti Crb (Cq4, 1:10). Chicken anti-Sr (1:20) was made against amino acids 707-1180 of the Sr protein fused to GST in the pGEX-KG expression vector. The ~80kDa fusion protein was purified on Glutathione-agarose beads followed by elution with reduced glutathione. Antibodies against the GST-Sr fusion protein were produced in Chickens by Dr. Enav Bar-Shira (Department of Animal Sciences, Robert H. Smith Faculty of Agriculture Food and Environment, The Hebrew University of Jerusalem, Rehovot, Israel). The IgY antibody fraction was isolated from the egg yolk and cleaned on Glutathione-agarose beads to reduce background of anti-GST antibodies. Secondary antibodies for fluorescent staining were Cy3, Cy2, Cy5 or Alexa Fluor-647-conjugated anti-Mouse/Rabbit/Rat/Chicken/Guinea pig (1:100, Jackson Laboratories, Bar-Harbor, Maine, USA). Stained embryos were mounted in DAKO mounting medium (DAKO Cytomation, Denmark) and viewed using confocal microscopy (Axioskop and LSM 510, Zeiss).

For the analysis of reporter gene expression, images were analyzed using ImageJ software (http://rsb.info.nih.gov/ij/) as previously described (Preger-Ben Noon et al., 2018). Briefly, maximum projections of confocal stacks were assembled, and background was subtracted using a 50-pixel rolling-ball radius. Then, we manually segmented visible nuclei of cap and scolopale cells from LCh5 of abdominal segments A2 and measured the fluorescence mean intensities of each nucleus. Statistical analyses and graphing were performed using GraphPad Prism version 8, GraphPad Software, La Jolla California USA, www.graphpad.com.

### Yeast-one-hybrid screen

The screen was performed by Hybergenics Services on a Drosophila whole embryo library (ULTImate Y1H screen *vs* Drosophila Whole Embryo RP2 0-12+12-24hr), using the *dei*^*ChO-389*^ sequence as a bait. Screening was performed using the Aureobasidin A selection system. 146 clones that were found to grow on the selective medium containing 400 ng/ml of the yeast antibiotic agent Aureobasidin A were sequenced (Table S1).

### Protein purification

*D-Pax2-HD* and *Pros-S-HD* expression plasmids were a kind gift from Brian Gebelein and Tiffany Cook (Cook et al., 2003; Li-Kroeger et al., 2012). Proteins were purified from *E.coli* (BL21) as described previously (Uhl et al., 2010) with the following modifications. Protein expression was induced at 37°C using 0.1mM IPTG for 4 hours (Pros-S/L-HD) or 0.4mM IPTG for 4 hours (D-Pax2-HD).

Pros-S-HD: After induction, bacterial pellet was resuspended in PBS supplemented with complete protease inhibitor mix (Roche) and lysed on ice using sonication (10 cycles of 30 sec on/off). GST-tagged Pros-S-HD in soluble fraction was purified using Glutathione-Agarose beads (Sigma). The bound proteins were eluted in elution buffer (50 mM Tris, pH 8, 10 mM reduced Glutathione).

D-Pax2-HD: The bacterial pellet was resuspended in 20 mM Tris, pH 7.5 supplemented with protease inhibitor. His-tagged proteins in soluble fraction were purified using cOmplete His-Tag Purification Columns (Roche). The columns were washed with 10 column volumes of wash buffer 1 (20 mM Tris, pH7.5, 300 mM NaCl, 50 mM NaH_2_PO_4_ pH 8.0), 2 column volumes of wash buffer 2 (20 mM Tris, pH7.5, 300 mM NaCl, 5 mM DTT, 10 mM Imidazole), and bound proteins were eluted in the same buffer supplemented with 250 mM Imidazole.

All samples were dialyzed against 500 ml of dialysis buffer (20 mM HEPES, pH 7.9, 200 mM NaCl, 10% Glycerol, 2 mM MgCl) for 18 hours at 4°C. Protein concentrations were measured with NanoDrop and confirmed by SDS-PAGE and Coomassie blue analysis.

### Electromobility shift assay (EMSA)

DNA probes were generated by annealing 5’ IRDye®700 labeled forward oligonucleotides with unlabeled reverse oligonucleotides (Integrated DNA Technologies) to a final concentration of 5 μM in PNK buffer (New England Biolabs). One hundred femtomoles of labeled IRDye®700 probes were used in a 20-μl binding reaction containing 10 mM Tris, pH 7.5; 50 mM NaCl; 1 mM MgCl2; 4% glycerol; 0.5 mM DTT; 0.5 mM EDTA; 50 μg/ml poly(dI–dC); 200 μg/ml of BSA and purified proteins (see Table SX for amount of each protein used). The binding reactions were incubated at room temperature for 30 min, and run on a native 4% polyacrylamide gel for 1.5 hours at 180 V. For competition assays, the appropriate amount of cold competitor was added with the IRDye®700-labeled probe prior to the incubation. The polyacrylamide gel cassettes were imaged using an Odyssey Infrared Imaging System and image analysis was performed using ImageQuant 5.1 software. All experiments were performed at least three times.

### Locomotion assays

Larvae used in the locomotion assays were collected from 8-12-hour egg collections that aged at 24°C until reaching the wandering 3^rd^ instar stage (115-140 hours). 25-30 larvae of each genotype were individually transferred to a fresh 2% agar 10cm plate, prewarmed to 24°C. Larvae were let to adjust for 30 seconds prior to 2-minute recording at a rate of 30 frames per second. The wild-type larvae often exited the filmed arena before the completion of the full 2 min recording time. Larval locomotion was recorded using a Dino-Lite digital microscope placed above the plate. We used VideoPad software to convert Dino-Lite files into Tiff files. ImageJ and FIMTrack (Risse et al., 2017) tracking software were used for following larval (center of mass) movements and body angle.

## Acknowledgements

We wish to thank Joshua Kavaler, Tiffany Cook, Anna Jazwinska, Manzur Bhat, the Bloomington Stock Center and the Developmental Studies Hybridoma Bank at the University of Iowa for sending us antibodies and fly strains. We are very grateful to Enav Bar-Shira for her kind help in generating the chicken anti-Sr antibodies and to Abeer Hassan for helping us with the analysis of the locomotion assays’ data. This work was supported by a grant (No. 674/17) to AS from The Israel Science Foundation.

