## Supplemental data for "*delilah, prospero* and *D-Pax2* constitute a gene regulatory network essential for the development of functional proprioceptors"

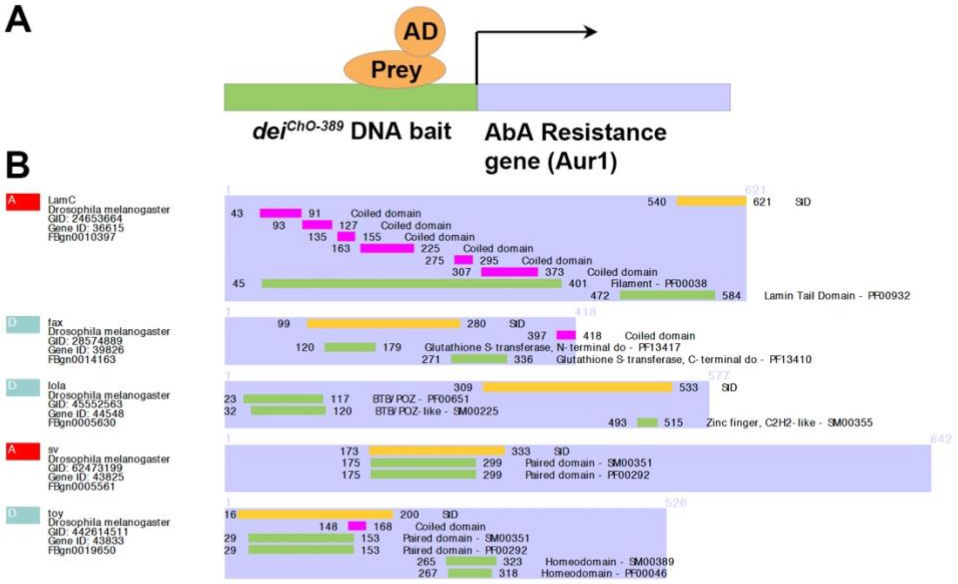


**Figure S1: Identifying potential direct regulators of dei in the ChO using a YiH screen.**

(A) Schematic representation of the screening system that used the dei$^{ChO-389}$ enhancer as a bait and the Aurobasidin A selection system to screen an embryonic cDNA library for preys that bind to the bait (Hybergenics Services). (B) A DomSight graph showing the top candidates identified in the screen in very high confidence in the interaction (red) or moderate confidence (green). The orange bars mark the SID fragment (selected interaction domain), which is the amino acid sequences shared by all prey.

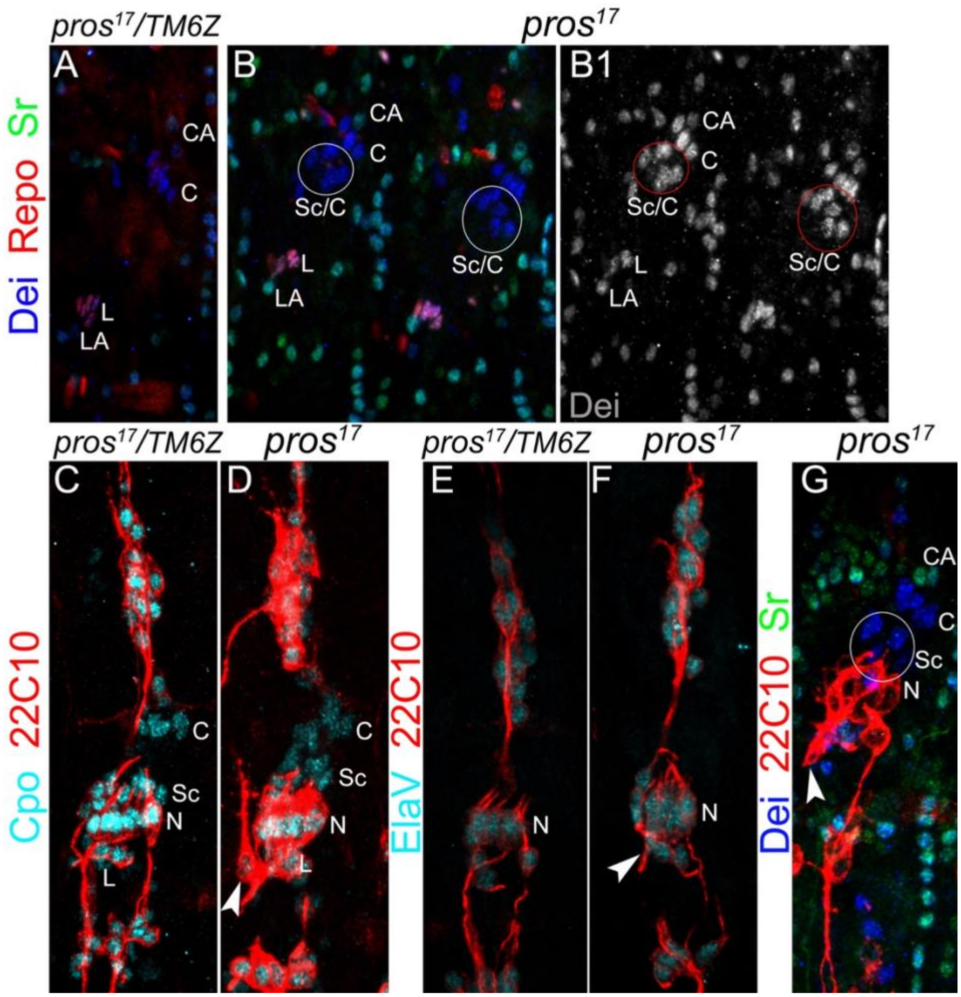


**Figure S2: The number of neurons and ligament cells remain normal in *pros* mutant embryos.**

(A-B) Representative segments of stage 16 *wt* (A) and *pros*$^{17}$ mutant (B) embryos stained for Dei (blue), Repo (red) and Sr (green). Repo staining shows the normal number of ligament cells in the mutant. (C-D) Representative LCh5 organs of *wt* (C) and *pros*$^{17}$ mutant (D) embryos stained for CPO (cyan) and 22C10 (anti Futsch, red). CPO is expressed in all of the LCh5 cells. (E-F) Representative LCh5 organs of *wt* (E) and *pros*$^{17}$ mutant (F) embryos stained for the neuronal markers Elav (cyan) and 22C10 (red). The data shown in C-F demonstrate that the number of neurons remains unaltered in *pros* mutant embryos. (G) An LCh5 of *pros* mutant embryo stained for the neuronal marker 22C10 (anti-Futsch), Dei and Sr. The arrowhead points to the typical axonal pathfinding defect of *pros* mutant embryos.

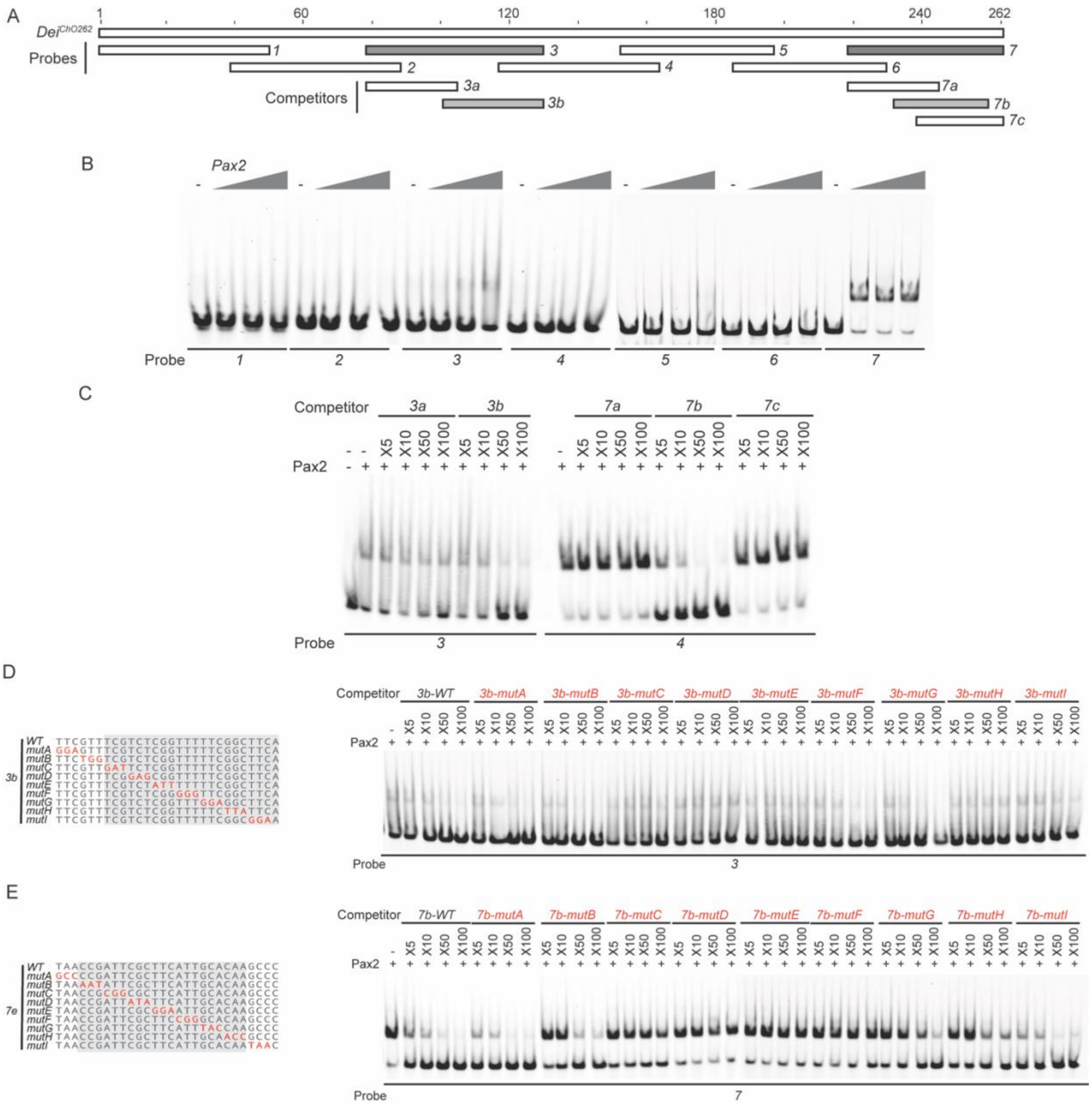


**Figure S3**. **Identification of the regions in the *dei*$^{ChO262}$ enhancer that bind D-Pax2/Sv *in vitro.***

(A) A schematic representation of the regions tested for their ability to bind D-Pax2/Sv in EMSAs. In gray are fragments that show specific binding. (B) EMSA scan for D-Pax2/Sv binding sites in *dei*$^{ChO262}$ using the oligos marked in (A). Two sub-regions of *dei*$^{ChO262}$ (fragments 3 and 7) bind to

D-Pax2/Sv. (C-E) Dissection of the D-Pax2/Sv binding sites through competition assays. EMSAs were performed in the presence of increasing concentrations of unlabeled oligos (competitors). (C) Sub-fragments 3b and 7b unlabeled probes completely compete the binding of fragment 3 and 7, respectively, to D-Pax2/Sv. (D-E) On the left, alignment of the mutated competitors used in the competition EMSAs shown on the right. The nucleotides that comprise the D-Pax2/Sv binding sites are marked in gray.

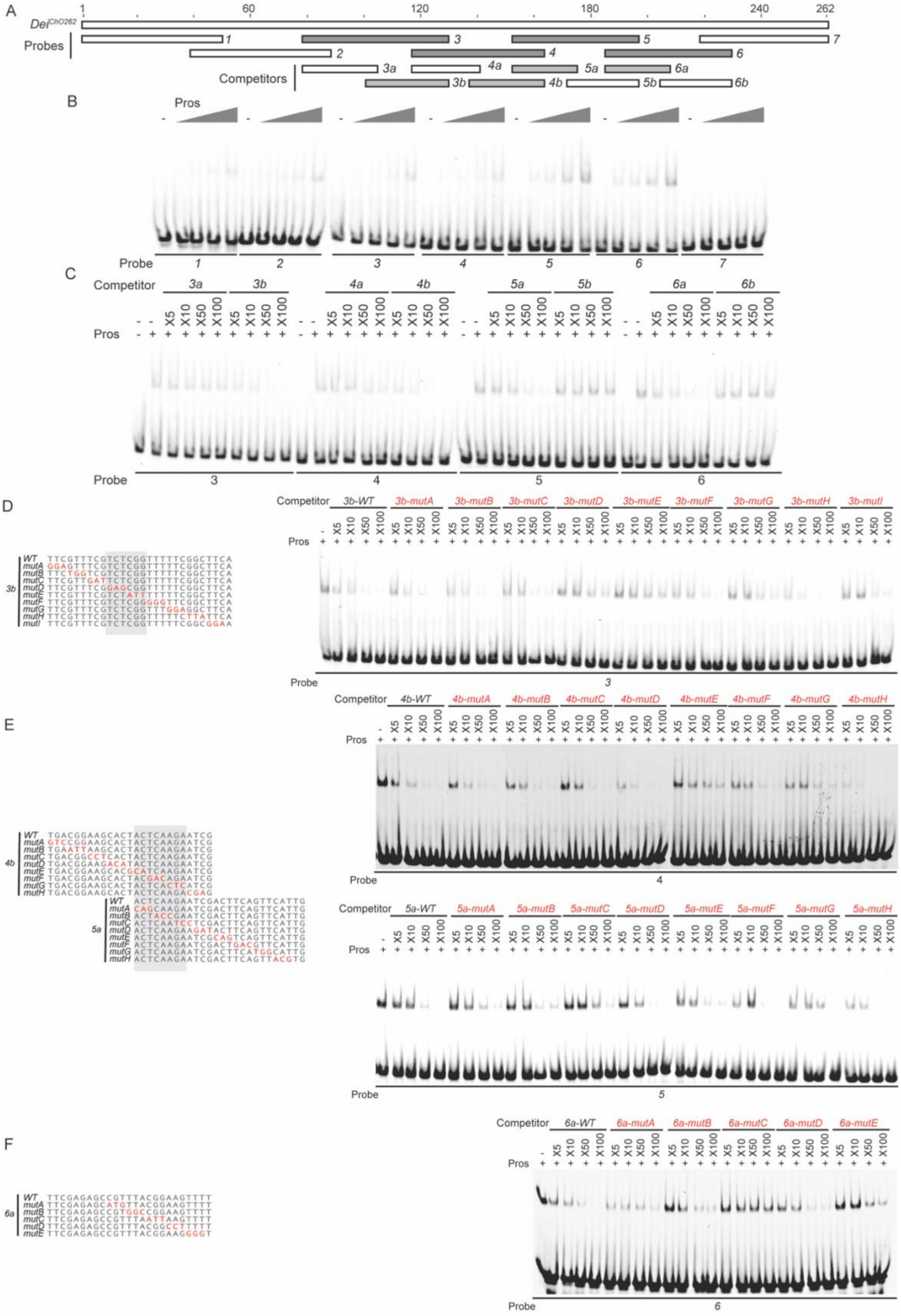

A
1
60
120
180
240
262
Dei^ChO262
Probes
1
2
3
4
5
6
7
Competitors
3a
3b
4a
4b
5a
5b
6a
6b
B
Pros
Probe
C
Competitor
X5
X10
X50
X100
Pros
D
3b-WT
3b-mutA
3b-mutB
3b-mutC
3b-mutD
3b-mutE
3b-mutF
3b-mutG
3b-mutH
3b-mutI
WT
mutA
mutB
mutC
mutD
mutE
mutF
mutG
mutH
mutI
E
4b-WT
4b-mutA
4b-mutB
4b-mutC
4b-mutD
4b-mutE
4b-mutF
4b-mutG
4b-mutH
5a-WT
5a-mutA
5a-mutB
5a-mutC
5a-mutD
5a-mutE
5a-mutF
5a-mutG
5a-mutH
F
6a-WT
6a-mutA
6a-mutB
6a-mutC
6a-mutD
6a-mutE

**Figure S4**. **Identification of the regions in the *dei*$^{ChO262}$ enhancer that bind Pros *in vitro.***

(A) A schematic representation of the regions tested for their ability to bind Pros in EMSAs. In gray are fragments that show specific binding (stable upon increase of non-specific and specific cometitors). (B) EMSA scan for Pros binding sites in *dei*$^{ChO262}$ using the oligos marked in (A). Four sub-regions of *dei*$^{ChO262}$ (fragments 3, 4,5 and 6) bind specifically to Pros. (C-F) Dissection of the Pros binding sites through competition assays. EMSAs were performed in the presence of increasing concentrations of unlabeled oligos (competitors). (C) Sub-fragments 3b, 4b, 5a and 6a unlabeled probes completely compete the binding of fragment 3, 4, 5, and 6, respectively, to Pros. (D-F) On the left, alignment of the mutated competitors used in the competition EMSAs shown on the right. The nucleotides that comprise the Pros binding sites are marked in gray.
