## Supplementary material for "*delilah, prospero* and *D-Pax2* constitute a gene regulatory network essential for the development of functional proprioceptors": Legends to Table S1 and Videos S1-6

**Table S1: Results of the 1YH screen**

This Table summarizes the sequencing data of 146 positive clones identified in the 1YH screen.

**Videos S1-6: Larval locomotion**. Videos S1-S6 show representative video recording of larval locomotion. Videos S1-S2 show *wt* (*Canton-S*) larvae. Videos S3-S4 show *dei^KO-mCherry^* larvae. Videos S5-S6 show *dei^ΔChO^* larvae.
